# Single-Cell RNA-Seq Reveals Naïve B cells Associated with Better Prognosis of HCC

**DOI:** 10.1101/731935

**Authors:** Jian He, Yingxin Lin, Xianbin Su, Qing Luo, Shila Ghazanfar, Jean Y H Yang, Ze-guang Han

**Author notes:** Equal contribution (Jian He, Yingxin Lin).

## Abstract

Hepatocellular Carcinoma (HCC) is a type of malignant solid tumor, causing high morbidity and mortality around the world and the major portion of HCC patients is from China. Cancer immunotherapies have shown some clinical responses in treating some types of cancer but did not shown significant efficiency in HCC treatment. This in part due to the impact of immune cells in the tumor microenvironment. It is commonly believed that HCC is a heterogeneous solid tumor and the microenvironment of HCC plays an important role in tumorgenesis and development. Currently, the residents of the microenvironment of HCC is not well-defined and clarification, especially the immune cells, which we believe that paly pivotal roles in tumorgenesis and development. To depict the landscape of the composition, lineage and functional states of the immune cells in HCC, we performed single-cell RNA sequencing on Diethylnitrosamine (DEN)-induced mouse HCC model. We observed heterogeneity within the immune and hepatocytes both in the precancerous condition of tumorigenesis and cancerous condition of HCC. In this study we found that the disease-associated changes appeared early in pathological progression and were highly cell-type specific. Specific subsets of T and B cells preferentially enriched in HCC, and we identified signature genes for each subset. Additionally, we mapped this group of specific cells to the human TCGA database. We found a cluster of naïve B cells characterized by high expression of CD38 associated with better prognosis of human HCC. Our study demonstrates signaling interaction map based on receptor-ligand bonding on the single-cell level could broaden our comprehending of cellular networks in varies status. Our finding provides a new approach for patient stratification and will help further understand the functional states, dynamics and signaling interaction of B cells in hepatocellular carcinoma, and may provide a novel insight and therapeutics for the HCC.

## Introduction

More than 50% of the worldwide cases of hepatocellular carcinoma (HCC) occur in China, and this malignancy currently solid tumor represents the country’s second leading cause of cancer death [1]. Recent studies have indicated that despite the lack of uniform efficacies among patients in different cancer types, cancer immunotherapies have dramatic impact on the oncological treatment outcomes in colon, lung, breast cancers and melanoma [2, 3]. In contrast, there has been a lack of clinical success in HCC therapies [4] due to the strong dependence on the amount and status of the immune cells in the HCC compliant tumor microenvironment (TME). Recent advances in this area has demonstrated a number of factors that correlate with clinical responses for check point therapies [5]. Unfortunately, these results haven’t been conclusive and have not been further developed to be applied as clinical biomarkers. It has been demonstrated that the hepatic microenvironment epigenetically shapes lineage commitment in mosaic mouse models of liver tumorigenesis by Lars Zender group [6]. To this end, detailed understanding of various immune cells in the community is crucial for the development of effective HCC immunotherapies and the identification of novel biomarkers. In particular, it is also pivotal to explore the composition and status of the immune cells during tumorigenesis. T cells and B cells are the most abundant and best-characterized population in the TME of solid tumors. Studies demonstrated that HCC tumor microenvironment (TME) harbor a significant level of T cells [7], while those TILs are incapable of killing tumor cells.

The ecosystem of liver tissue include hepatocyte, endothelial, fibroblasts, epithelial, immune cells and many other diverse cell lineages. Tumor formation requires the coordinated function and crosstalk between specific cell types in this distinct ecosystem. Immune cells migrate from hematopoietic organs to the liver and establish an active immune niche that interacts with stromal cells, influencing differentiation, tumorigenesis and development. At the same time, development of the cells in liver into specialized cells is a highly regulated process. Within a specific microenvironment, dominant signal transmitting cells determine the cellular composition and characteristics by secretion of protein signal. Therefore, The TME of HCC could be an appealing model to characterize the dysregulation and cross talk of various cells.

Single-cell RNA sequencing (scRNA-seq) provides an efficient method to study the cellular heterogeneity of the HCC, by profiling tens of thousands of individual cells [8, 9] simultaneously in the highly complex TME. Traditional bulk-tissue-level resolution may mask the complexity of alterations across cells and within cell groups, especially for less abundant cell types [10]. Potential changes in cell composition during HCC tumorgenesis will also confound the distinction between composition and changes in activity in a given cell type. Thus scRNA-seq profiling of cells enables the decomposition of cell types experimentally and ascertain which of these features may predict or explain clinical responses to anticancer agents. Recently study in Melanoma has shown that this approach when applied to characteristic cells from melanoma patients was able to reveal the exhaustion signature of T cell [12].

Most human liver cancer studies are cross-sectional at specific given time point and thus unable to provide good characterization of tumorgenesis, such as the dynamics, status and transformation of the cancer development. To begin addressing some of these limitation, we design a study based on DEN induced mice that enable the comparison between the early and late stage of cancer development. This is an initial attempt to capture the dynamics of the cell types and status during the tumorigenesis and development. Note that, DEN model is a well-established chemical induced model that could mimic human HCC [13]. The DEN induction is the most commonly used method in chemically induced liver cancer models, because DEN is metabolized in the liver, it is most effective at inducing liver tumors.

In this study, the single-cell RNA-seq examine >8,000 cells isolated from DEN induced mice enabling us to simultaneously study their transcriptomes. We identified 36 unique cell subsets with distinct tissue distribution patterns, status and interactions. This large-scale transcriptome data of immune cells, especially the transition of status of T cells and B cells and the correlation between immune and hepatocytes can be used as a profitable resource for profiling the characteristics of immune cells and for providing novel insight and approach into HCC prevention and treatment. Moreover, the complex interplay within and across cell types, further contributes to the difficulty in interpreting tissue-resolution signatures of HCC. Our single-cell transcriptomic resource provides a blueprint for interrogating the molecular and cellular basis of HCC.

## Results

### Landscape of the cell composition and characterization during DEN-induced Liver cancer

To depict the landscape of the composition, lineage and functional states of the immune cells in hepatocellular carcinoma (HCC) tumorgenesis and development, we systematically study the composition and characteristics of different immune and non-immune cell types and status for HCC tumorigenesis by perform single-cell profiling of tumorigenesis and tumor microenvironment.

We collected DEN-induced 6-month and 16-month mouse liver tissue, and we observed that 6 months were pre-cancerous (or early-stage HCC pathology) and 16 months were cancer (or late-stage HCC pathology) (SI-Fig 1). We can see a large group of inflammatory cells were recruited in the pre-cancer (or early-stage HCC pathology). The tumor tissue of the 16M cancer mouse were liquefactive necrosis at the same time, the adjacent tissue contained numerous immune cells (Supplementary Information-Fig 1, SI-Tab 1). In order to avoid selective tissue dissociation procedures, we use a permissive tissue dissociation protocol, resulting in a comprehensive repertoire of the immune and non-immune cells located in the liver.

Unsupervised clustering analysis with the most highly variable genes (see Methods) reveal distinct patterns of cell clustering at two levels and differential expression analysis and differential variation analysis were used to facilitate with cell type annotation. We are able to annotate the cluster into 10 main groups: T cell, B cell, Neutrophil cell, Hepatocyte, Dendritic Cell, Endothelial, Epithelial, Fibroblast, Erythrocyte (Fig 1a, SI-Fig 2).

**Fig 1.**
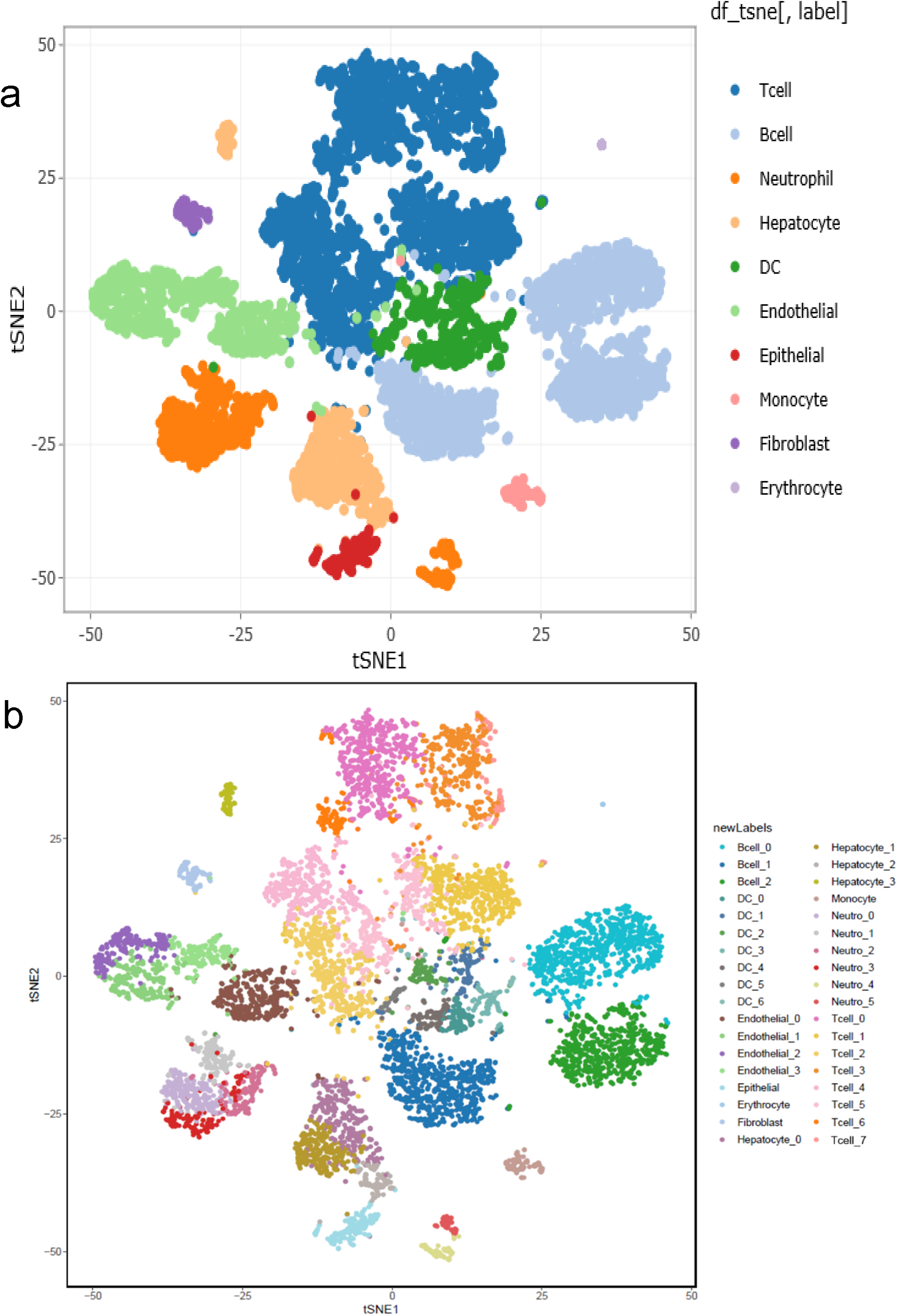
t-SNE map displaying 8,872 cells. (a) Over all profile. (b) Subsets of each cell types.

This indicates that in the HCC tumorgenesis and development, there was a large amount of immune cell (78.9%) infiltration in addition to liver cells. The presence of a small group of erythrocyte (0.2%) indicate that maybe some blood residual in the liver. Furthermore, we found that the cell type compositions are more similar for samples within the same time point than between early and late stage tumorigenesis. In particular, the proportions of T cell and DC significantly increase (p-value from scDC < 0.001, see Method) in the late samples (Fig 2). The proportions of Neutrophil cell, Hepatocyte, Endothelial, Epithelial and Fibroblast decreased in the late stage.

**Fig 2.**
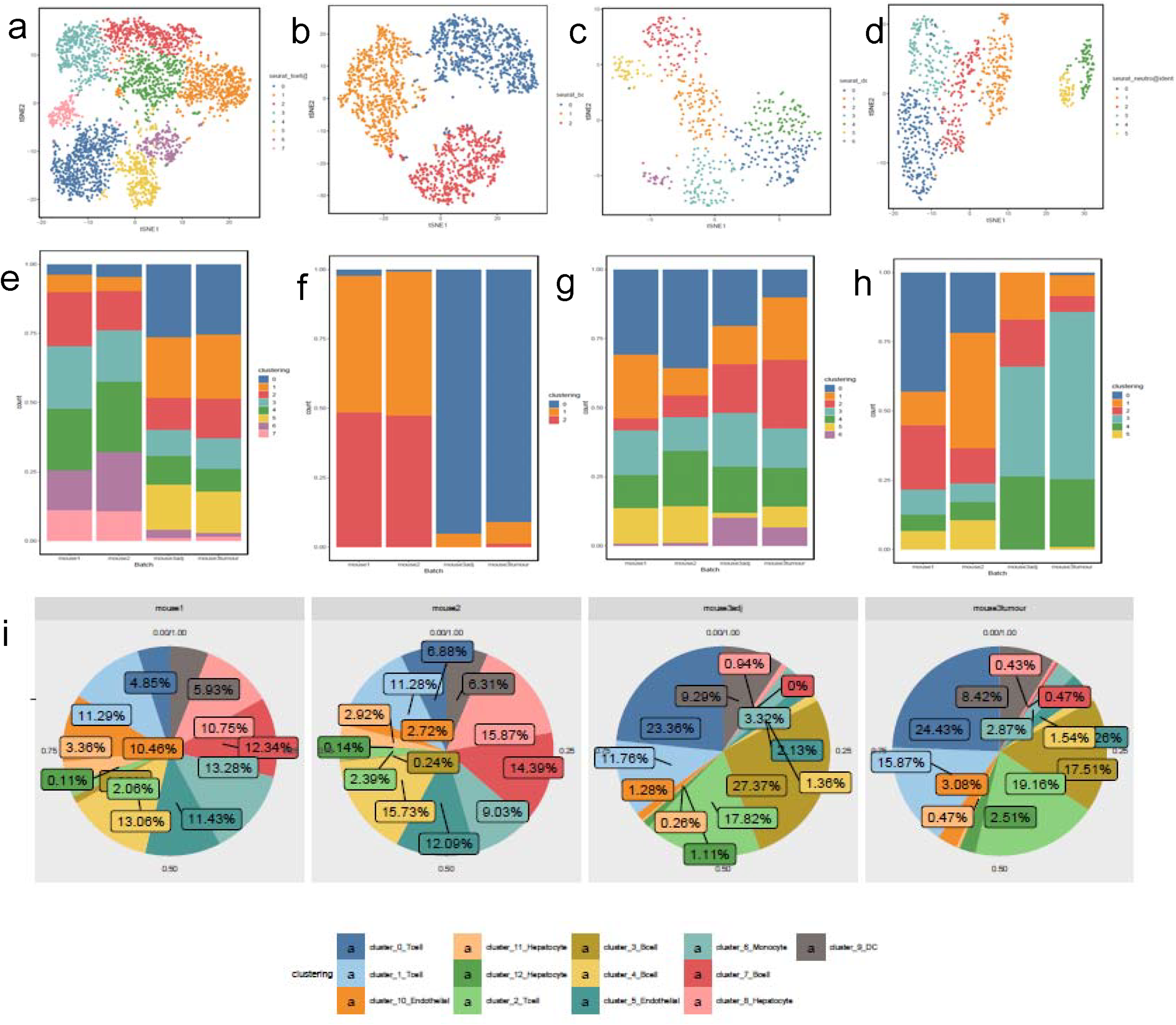
Tissue micro environment affects the diversity of immune phenotypic states. (a, e) T cells, (b, f) B cells, (c, g) DC, (d, h) Neutrophil, (i) cell type composition by tissue.

We performed unsupervised clustering of each cell types to discover further structure using the spectral clustering method implemented in Seurat [14]. This refined result generated a total of 36 subgroups (Fig 1b, Method) with T cells having the largest number of sub-groups (8 subgroups; Fig 1b, Fig 2a, 2g) illustrating its diversity characteristics. In this study, DC were divided into 6 subsets may indicated various responses under different stimulations. It is interesting to note that the cell type composition is similar between tumor and adjacent tissue at the ‘main’ level but DC, Neutrophil, Endothelial subgroups are distinctly different at the subgroup level. This analysis identified the main immune cell types in both T cell (8 sub types), B cell (3 sub-types) and other immune cells. T cells were the main immune cell population in the HCC TME, with a mean of 36.4% across samples. The mean frequencies of B cells (25.9%), Neutrophil (8.6%), Hepatocyte (7.9%), DC (6.7%), Endo (10.1%), Epi (1.8%), Monocyte (1.3%), Fibroblast (1.1%) and Erythrocyte (0.2%), respectively (Fig 2 and SI-Tab 2) suggesting beside the original resident, a large number of immune cells are resident in this pathological environment. It is interesting that the total number of immune cells is as high as 70%, and the total number of T cells and B cells has exceeded the half of the total number of cells. This novel discovery demonstrates that studying immune cells is the vital for us to understand the process of tumor development. This implying that if we could depict the tumor immune microenvironment is, to some extent, equivalent to research and understanding the HCC pathogenesis. As expected, we observed that the immune cell composition and characteristics changes during tumorigenesis (between the early and late time point) as clearly observed in Figure 2.

### scRNA-seq profiles reveals the presence of malignant liver cells in precancer samples and the Afp progenitor marker in late stage samples

The pattern of hepatocyte shifts tempestuously not only from early to late stage but also between tumor and adjunct. A group of hepatocyte characterized by Afp+ subsets is accumulate in late stage, especially in tumor. These findings suggest that hepatocyte could transfer from Afp− to Afp+ in tumorgenesis and Afp+ hepatocyte is tumor-resident hepatocyte might be release cytokine to immune cells (Fig 3).

**Fig 3.**
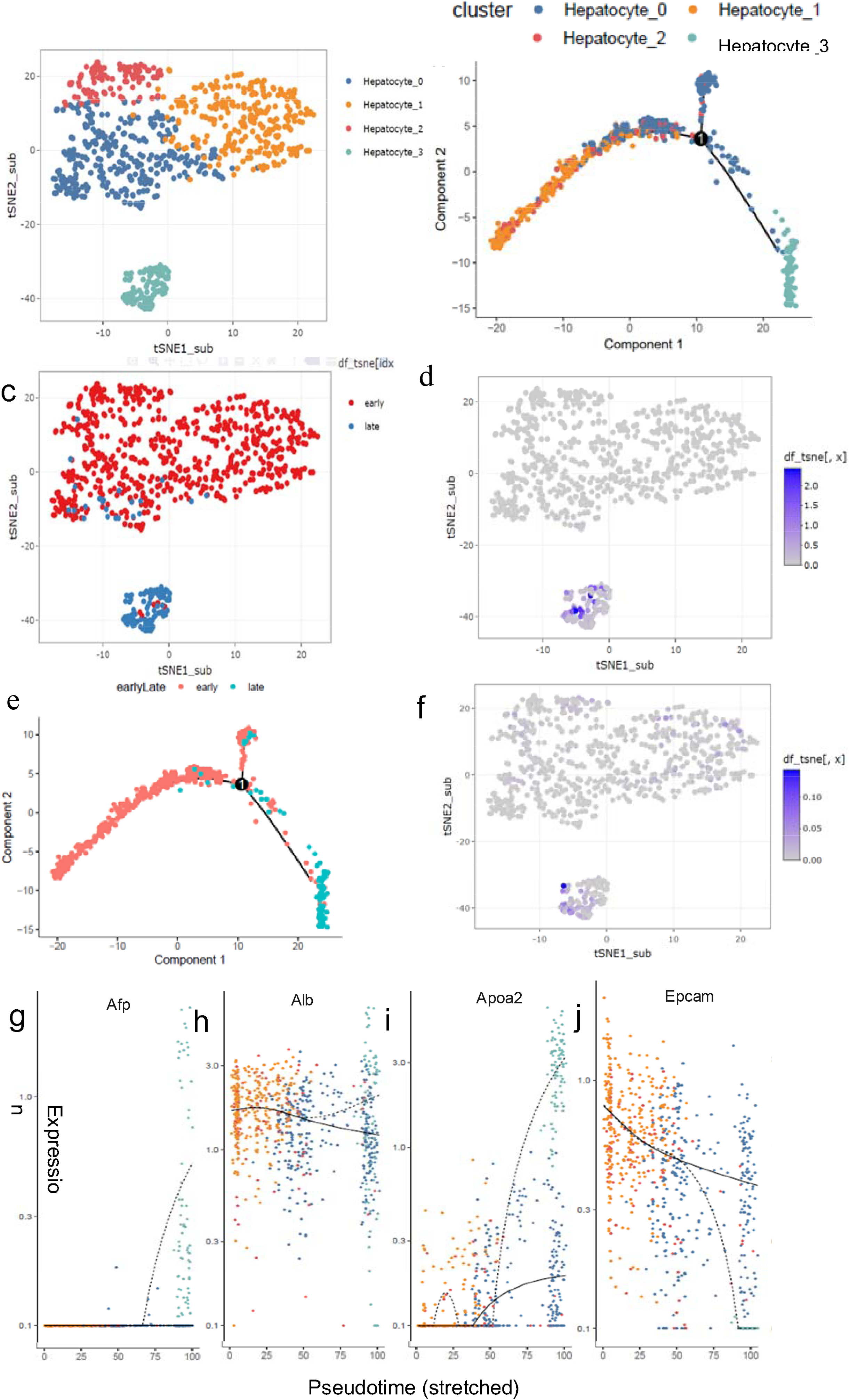
Single-cell transcriptome reveals malignant show up in precancer. (a) tSNE of the subclusters of hepatocyte, (b) pseudo-trajectory of hepatocytes, (c) early and late stage of hepatocytes, (d) Afp expression patterns in hepatocytes, (e) cell type transition from early to late stage, (f) Gpc3 expression patterns in hepatocytes,(g-j) Pseudotime of Afp, Alb, Apoa2, Epcam.

There were three subsets of liver cells during precancer stage and the response patterns were different, whereas it was more uniform in cancer stage, on the other side, there also many Afp− liver cells in the cancer stage. All these indicate that in the inflammation environment the liver cell demonstrated high heterogeneity. GPC3 also shares same pattern. In previous study based on bulk sequencing, we believe that of the Afp is negative in the precancer stage. In our study, we demonstrated that some cells in the early stage could also express Afp at a single cell resolution. This might be an indication of early cancer screening. We also found that the liver stem cell marker Met is highly expression in the early stage while Thy-1 does not demonstrate significant difference between the two stages.

### Focus examination of T and B cells immune cells suggests differential cell type compositions between early and late stage tumorigenesis

Detailed examinations of T and B cells suggests a further eight T-cell and three B-cells subgroups. T cells include seven clusters for CD8+T and one cluster for CD4+T cells, each with its unique signature genes [15-18] (Figures 4, Supplementary Information-Table 2). Differential composition analysis reveals different cell type proportion of T cells subgroups in early and late stage tumorigenesis. In particular, by fitting a fixed effect generalized linear model (GLM), we found that the proportions of Exhausted T cell (T0) and Irf8+T (T5) increase in the late stage tumorigenesis, with significant interaction effect of stage and subgroup of the model (p<0.001) MAIT (T1) also increase. Naïve T cell (T3), NKT (T4), Cd4+T (T6) decrease in late stage (Fig 2 a, i, Fig 4 a, b and SI-Tab 3).

**Fig 4.**
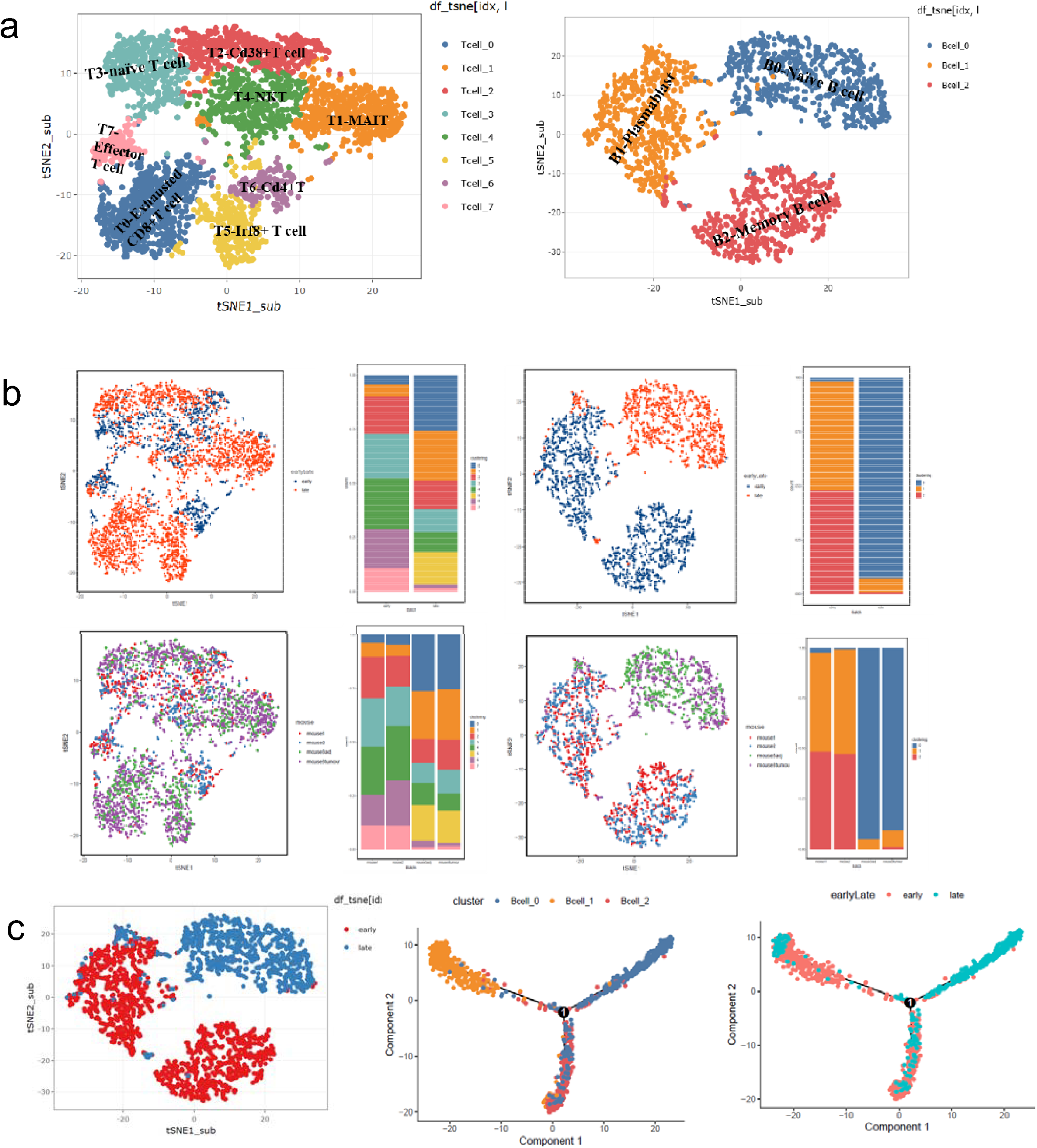
tSNE of the subclusters of T cells and B cells (a) and of early and late stage (b) and pseudotime of B cells.

Similarly, there is a clear increase in the proportion of Naïve B cell (B0) from the early to late stage tumorigenesis, while the proportions of Plasmablast (B1) and Memory B cell (B2) dramatically decrease in the late stage, confirming by the GLM analysis with significant interaction effects between B cell subgroups and stages (p<0.001). This dramatic changing of B cells suggests tumor microenvironment may inhibit maturation of naïve B cells (Fig 4c, Supplementary Information-Table 3).

### Diverse phenotype among PD-1+ T cells within tumorigenesis

We observe that intra-tumoral T cell clusters are characterized by diverse patterns of environmental signatures. Our findings suggest that tumor-resident T cells might be exposed to varying degrees of inflammation, hypoxia, and nutrient deprivation. While many of these responses (e.g., activation or hypoxia) individually represent phenotypic continuums, their combinations may result in more discrete states. Figure 5 shows that regulatory T cells (T6) were mostly observed among the CD4+ subset and were defined by the co-expression of CD25 and Foxp3. PD-1+cells were mainly observed in CD8+ subsets, especially in Exhausted CD8+T cell (T0) and naïve T cell (T3). The subset T0, characterized by the highest level of PD-1 expression among CD8+cells, was also positive for other co-inhibitory receptor HAVCR2, the activation markers CD38 and the co-stimulatory receptors such as TNFRSF9 and ICOS. This phenotype was associated with exhausted T cells and anti-PD-1 treatment response [19, 20]. Subsets with similar levels of PD-1 differed in expression of activation markers and co-stimulatory receptors. The cluster T3 had the same expression pattern as T0 but the activation markers and co-stimulatory markers were expressed at lower levels. The other PD-1+clusters (T2, T5 and T7) were characterized by the absence of HAVCR2 expression and by heterogeneity in expression of markers such as TNFRSF9, CTLA-4 and CD38. Unlike Cd4+ Regulatory T cells (T6) presented in the early stage, PD-1+ clusters were mainly present in late stage, but both of them did not show significant difference between tumor and adjacent tissue (Fig 5). All of the results above demonstrated a diversity phenotypic among PD-1+ T cells present in tumorigenesis.

**Fig 5.**
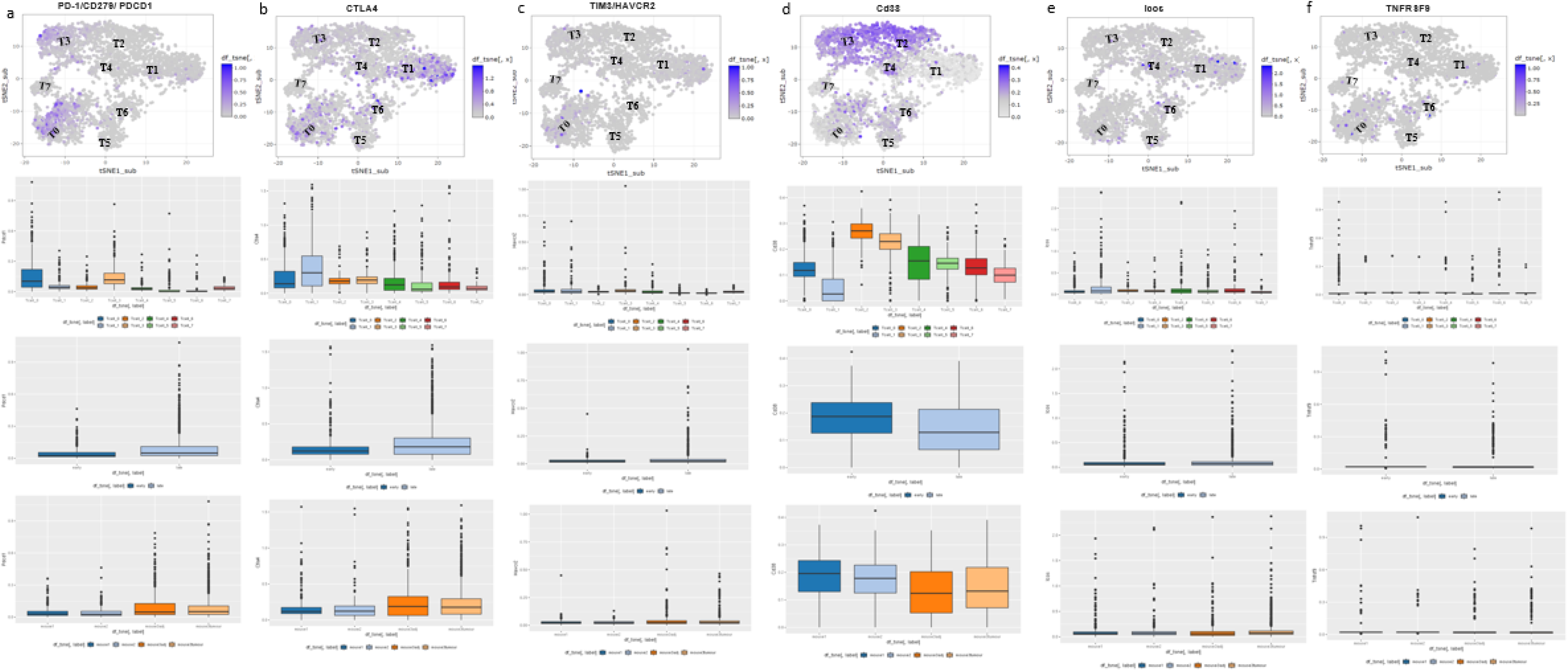
T Cell Characterized by immunosuppression associated markers.

### Analysis of cell-cell communication facilitate prediction of signaling interaction map at the single cell resolution

To understand the interactions between various cells in the HCC microenvironment of the DEN-induced mice thoroughly, we employed a ligand-receptor interacting repository which accounts for their subunit architecture (www.CellPhoneDB.org). This repository comprehend ligand–receptor interaction by the diffusion of secreted molecules [21, 22]. We explored the expression levels of ligands and receptors of each cell type and calculate which ligand-receptor pairs exhibit significant cell-type specificity. This could predict interactions between cell populations via specific ligand–receptor binding and generate a potential cellular communication network in the HCC microenvironment. B0-naïve B cells, Hepatocyte0, all subtypes of the Endothelial and DC6 (Ccl17+DC), which are dominant in the late stage, harbored the highest “outgoing” signals. Hepatocyte3 (late stage Hepatocyte) and dysfunctional T cells-harbored the highest “incoming” signals (SI-Tab4). All subtypes of epithelial are harbored both “outgoing” and “incoming” signals. B0-naïve B cells could bind CCR2 which is on the dysfunctional T, by releasing cytokine CCL2, CCL24, CCL8, CCL11 and bind CXCR3 by releasing cytokine CCL20, in this way, naïve B cells could regulate T cells. Meanwhile, Hepatocyte3 (late stage Hepatocyte) could bind Igf1r which is on the B0-naïve B cells, by releasing cytokine Igf1, in this way, late stage Hepatocyte could regulate B0-naïve B cells. What’s interesting is that, naïve B cell binds Clta4 on all types of T cells except T7-effect T cell, by releasing CD86. Fig 6 and SI-Fig 3 showed an overview of selected ligand–receptor interactions, indicating that T cells are likely the “terminal”, receiving single from other cells, especially from endothelial, epithelial and hepatocyte. B cells are more likely as “originating”, they could release single to other cells, at the same time they can “self-regulation” (Supplementary Information-Table 4). All the above reveal regulation mechanisms of different cell types in HCC tumor microenvironment.

**Fig 6.**
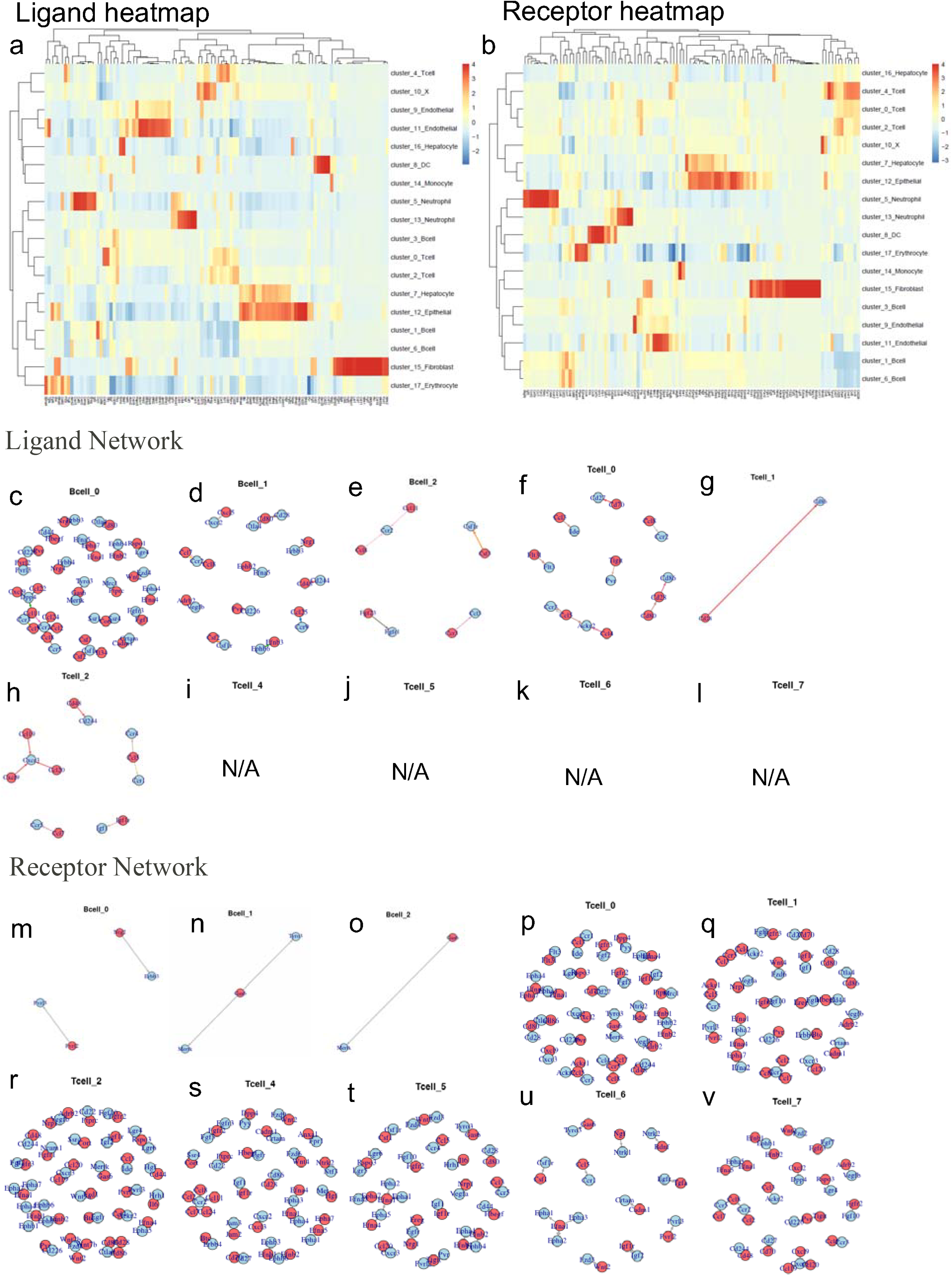
Cell communication predicted by CellPhoneDB. Ligand heatmap (a). Receptor heatmap (b). Ligand network of B cells and T cells (c-l). Receptor Network of B cells and T cells (m-v).

### Naïve B cell markers associated with better prognosis of HCC (Compare single cell data with TCGA data)

To compare the cell type proportion between the primary solid tumor samples and normal solid tissue samples of human, we used the single-cell RNA-seq data to decompose the cell types of the TCGA LIHC bulk RNA-seq data (see Method). We have found that statistically significant differences of proportion exist in several cell types (Fig 7), such as memory B cell (B2), Dendritic cell (DC2) and Afp+ hepatocyte (H3), suggesting scRNA-seq could demonstrate more detail information, and dig out the information that concealed in “average expressing” pattern. And also some clusters share same pattern with the bulk sequencing data (Red asterisk: Bulk data is consistent with scRNA-Seq data; Blueline: Bulk asterisk is different from scRNA-Seq data) tested by Wilcoxon Rank Sum test [22]. To obtain a global understanding of the relationships between all immune populations and clinical outcome, we applied a Cox proportional hazards model that included age, gender, tumor stage in addition with the proportion of certain cell type in the sample (Fig 7b). We observed correlation between the proportion of cell types with survival for immune cells. In particular, we found that the patients with higher proportion of Exhausted CD8+T cell (T0), MAIT (T1), naïve B cell (B0) have higher chance of survival, while lower proportion of Cd38+T cell (T2), naïve T cell (T3) and Irf8+T cell (T5) has higher survival rate.

**Fig 7.**
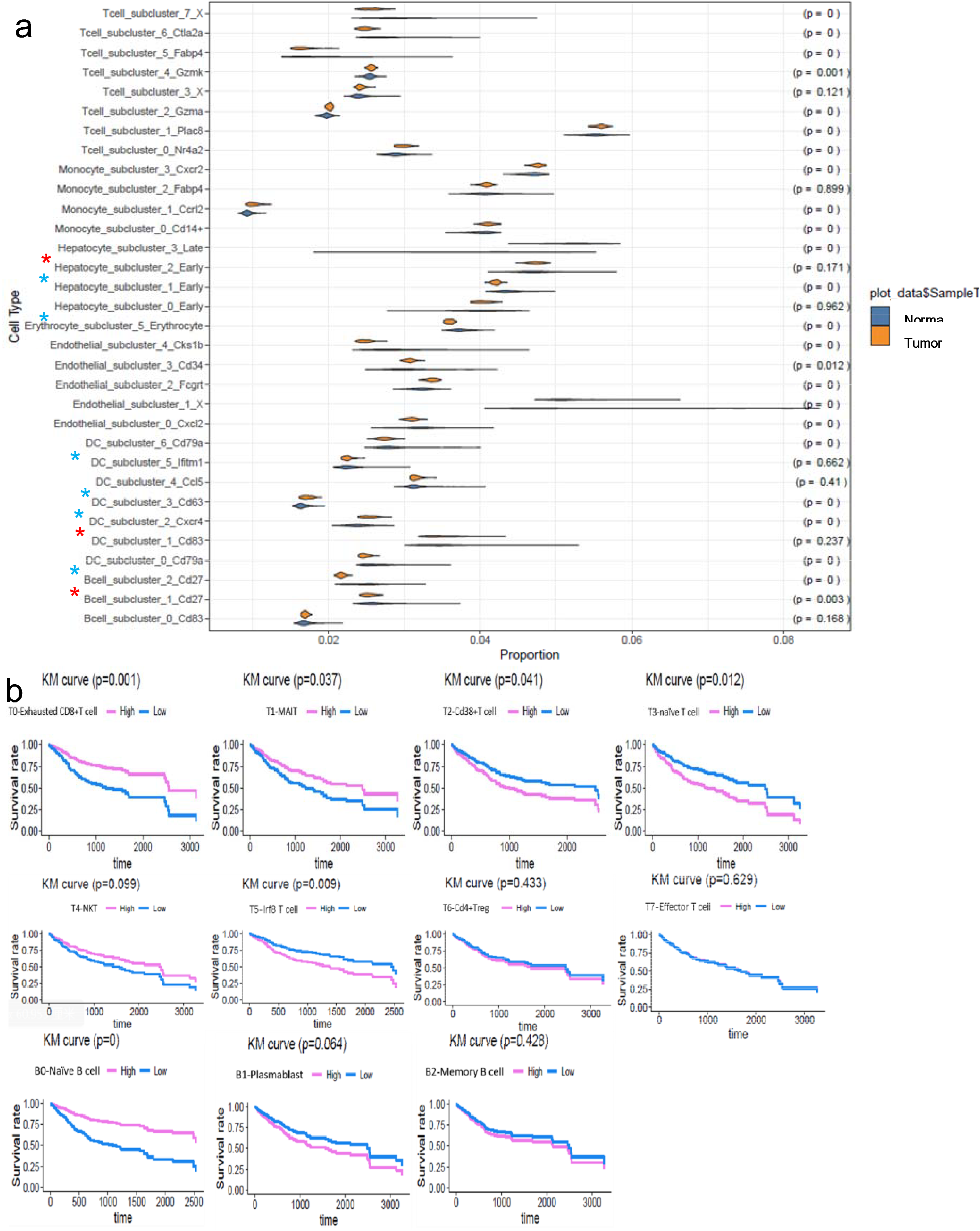
(a) Cell Type Decomposition with Bulk TCGA RNA-seq data, (b) and correlation between the proportion of cell types with survival rate.

## Discussion

ScRNA-seq provides an cutting edge method to study the cellular heterogeneity of the tumor microenvironment of many cancer types, by profiling the transcriptomics of thousands level of individual cells [8, 9]. Bulk-tissue-level resolution may mask the complexity of alterations across cells and within cell groups, especially for less abundant cell types [24]. Potential changes in cell composition during HCC tumorgenesis also confound the distinction between composition and activity changes in a given cell type. Moreover, the complex interplay between protective and damaging molecular processes, both within and across cell types, further contributes to the difficulty in interpreting tissue-resolution signatures of disease. In our study, the single cell sequencing data along with the transcriptome information for >8,000 individual cells provided a comprehensive resource for understanding the multi-dimensional characterization of cells in HHC tumor microenvironment, especially immune cells and hepatocyte. The pivotal role of T cells in anti-tumor responses has been widely studied and well established. However, the importance of B cells to the immune response to tumorigenesis and development are not well defined. With the development of the higher resolution technology and careful design study design, we are witnessing more and more evidence supporting a role for B cells in tumor immunology. Recently, it has been reported that the presence of tumor-infiltrating B lymphocytes has been linked to favorable clinical outcome in many types of cancers [25]. However, B cells represent a heterogeneous population with functionally distinct subsets, and the dynamic transition process among subtypes impacts tumor development. B cell–associated autoimmune responses have been commonly found in various tumor types. A wide range of autoantibodies is found to be as markers or as direct mediators of paraneoplastic syndromes [26], whereas autoimmune diseases are associated with increased or reduced risk of cancer is unclear. Recently, Joseph group demonstrated that cancer can trigger acquired humoral immunity, showing that the appearance of certain autoantibodies is part of a defensive (although eventually unsuccessful in some cases) immune response against a developing tumor [27]. Autoantibodies have attracted interest as potential biomarkers for early diagnosis or for their potential as prognostic indicators for many types of cancer [26].

It has been proved that the autoantibodies of tumor associated B cells has become potential biomarkers for early diagnosis or for their potential as prognostic indicators for many types of cancer. Although the mechanism through which autoantibody responses are triggered, particularly against commonly expressed proteins, such as glycolytic enzymes, is not well understood. The basic function of the immune system is to discriminate self from non-self. Since immune tolerance against self-antigens is a highly regulated process, the presence of autoantibodies may signal either breakdown of immune tolerance or that some self-antigens are not encountered by immune cells in nonpathologic states because of their intracellular turnover and limited accessibility in nonmalignant cells [28]. Self-antigens reactivity was not only a dominant characteristic in tumor-bearing mice samples but also in prediagnostic human samples [29]. The evolution in autoantibody reactivity during tumorgenesis and development further demonstrates the dynamics of the autoantibody response. Some cancer-associated autoantibodies are triggered by cellular proteins that are mutated, modified, or aberrantly expressed in tumor cells and hence are regarded as specific "immunological reporters" [30]. The concept that new antigenic epitopes may arise from protein processing has both diagnostic and therapeutic relevance [30].

The gene signatures of B-cell, especially B-TILs, have been associated with a favorable prognosis for several cancer types, including breast and ovarian [31, 32]. Schmidt group demonstrated that the presence of B-TILs has been linked to a favorable clinical outcome, including breast cancer and NSCLC [28, 33, 34]. These studies have corroborated that the presence of B cells, as derived from gene expression signatures, is linked to a favorable prognosis in breast cancer [32]. Concordantly, immunohistochemical staining of CD20 as a surrogate marker for B cells was found to have prognostic significance for B-TILs in breast cancer [28, 34]. And CD20 is the marker of naïve B cell. CD20 become low expression after naïve B cell matures. But the relationship between naïve B and prognosis of HCC has not been elicited before our study. Our group is the first one who established the naïve B cell is positively correlated with the prognosis in HCC.

B cells represent a heterogeneous population with functionally distinct subsets, contributing to both protumor as well as antitumor immune responses, and the balance among the subtypes may affect tumor development and behavior [35, 36].Previous study demonstrated that B cells in the tumor microenvironment is related to immune responses. But recent evidences show that B cells have the capacity to recognize antigens, regulate antigen processing and presentation, and mount and modulate T-cell and innate immune responses [28]. The tumor microenvironment contains a heterogeneous population of B cells with functionally distinct subsets, the collaboration of these responses contributing to both pro-immune as well as anti-immune function. In our study, the heterogeneity of B cells and the increase in late T cell suggesting B cells may regulate the quantity and status of T cells. Apart from the immune-regulatory function of antibody and antibody–antigen complexes, B cells can shape the functions of other immune cells by presenting antigens, providing co-stimulation, and secreting cytokines [37]. The antigen–antibody interactions in this study may, in part, on the extent of provide some evidence for dynamic changes in autoantibody reactivity of B cells contribute to tumor development and progression.

## METHOD DETAILS

### EXPERIMENTAL DETAILSDEN-induced HCC mouse model

C57BL/6 mice were purchased from Shanghai Laboratory Animal Center. Three mice which were DEN induced were enrolled in this study, including two DEN-induced 6 month and one DEN-induced 16M pathologically diagnosed with HCC. The mice were injected intraperitoneally 25 mg/kg DEN on the twelfth day after birth. The whole liver tissue of DEN-induced 6 month mouse and the paired fresh tumors and adjacent liver tissues of DEN-induced 16M mouse were obtained. The adjacent tissues were at least 2 cm from the matched tumor tissues. This study was approved by the Ethics Committee of Shanghai Jiao Tong University.

### Tissue dissociation and single cell suspension preparation

Fresh tissue samples were cut into approximately 1 mm^3^ pieces and gently triturated with a 20 mL syringe plunger on a 70 μ m cell strainer (BD) into the PBS (Invitrogen) with 10% FBS until uniform cell suspensions were obtained. The filter residue on the 70 μ m cell strainer were collected and digested for 20 min in 0.05% collagenase IV on the shaker in 37 □, then passed through a 40 μ m cell strainer into the cell suspension from the last step. The suspended cells were centrifuged at 200g for 5 min. After supernatant removal, the pelleted cells were suspended in red blood cell lysis buffer (Solarbio) and incubated on ice for 5 min to lyse red blood cells. After washing twice with 1 × PBS, the cell pellets were re-suspended in buffer.

### Hematoxylin & Eosin Staining

Warm the Slides on the slide warming table for 5-10 minutes. Repeat the treatment. Hydrate the sections by passing through the graded alcohol in decreasing order for 1-2 minutes (100%, 90%, 80%, 70%, 50%). Wash in tap water. Stain in hematoxylin for 3-5 minutes. Wash in running tap water for 5 minutes or less. Differentiate in 1% acid alcohol (1% HCl in 70% alcohol) for 5 minutes. (Check the Differentiation by using a microscope. The Nuclei should appear dark purple or Reddish blue and the tissues appears pale).Wash in running tap water until the sections are again blue by dipping in an alkaline solution followed by tap water wash. Stain in Eosin for 30s. Wash in tap water for 1-5 minutes. Dehydrate in increasing concentration of alcohols and clear in xylene. Observe under microscope.

### Library preparation

Cells were immediately counted using a hemocytometer and loaded in the 10x-Genomics Chromium. The 10x-Genomics v2 libraries were prepared as per the manufacturer’s instructions. Libraries were sequenced, on an Illumina HiSeq 2000 (paired-end; read 1: 26 cycles; i7 index: 8 cycles, i5 index: 0 cycles; read 2: 98 cycles).

## QUANTIFICATION AND STATISTICAL ANALYSIS

### Low level processing and filtering

All RNASeq libraries (pooled at equimolar concentration) were sequenced using Illumina HiSeq 2000 at a median sequencing depth of 120622 reads per single cell. Cellranger was used to perform demultiplexing, alignment, filtering, barcode counting, and UMI counting. Cells with less than 100 UMIs were discarded from the analysis. Seurat was used for normalization, dimension reduction, unsupervised clustering and visualization on the single cell transcriptomes. Monocle was used to predict pseudo time trajectories of specific cell clusters.

### Quality Control and Preprocessing

The cells were selected by performing empty Dropsto distinguish gene expression profiles that were significantly different to ambient expression profiles. In addition, we used k-means clustering on total expression and mitochondrial proportion to select the set of cells that are likely to be damaged (characterized by very high mitochondrial proportion). We further filtered the cells based on the total number of counts, total number of features, and mitochondrial proportion. After the quality control, 17493 genes and 8821 cells were used for downstream analysis. We then performed SAVER to impute the data to recover the gene expression from the dataset.

### Clustering and cell type identification

To identify and annotate the cell type from the data, we performed clustering analysis using Seurat on the first 20 principal components (PCs), which based on a shared nearest neighbored modularity optimization algorithm, with the resolution parameter as 0.8. Then we performed differential expression analysis using MAST and differential variability analysis using Bartlett test to identify the markers for each cluster, resulting to 10 main cell types in total. We further performed clustering analysis using Seurat, and differential expression analysis within each main cell type to identify sub clusters.

### Cell type differential composition analysis

To analysis the relationship of cell type proportion with mouse phenotypes, we performed differential compositional analysis (scDC) using the R package scdney (https://github.com/SydneyBioX/scdney). Bootstrap resampling was performed to capture the uncertainty of the main cell type proportion as well as sub cell type proportion within each main cell type. We then fitted the fixed effect generalized linear model (GLM) on the number of cells in each cell type by treating conditions of the sample as predictors on each bootstrap samples. The univariate Wald tests were used as significant tests on the pooled parameters from the GLM models.

### Trajectory analysis

To investigate the hepatocyte cell development trajectory during tumorgenesis and its development, we reconstructed the pseudo-time cell trajectory using monocle2, using the differential expressed genes between sub clusters of hepatocytes as ordering genes and DDRTree (discriminative dimensionality reduction tree) as dimension reduction method.

### Cellular communication analysis

To investigate the cell-cell communication between different cell types, we performed a similar analysis proposed by Vento-Tormo et al. by analysis the ligand receptor pairs provided in on CellPhoneDB. To identify the significant cell-cell interaction, for each ligand-receptor pair, we performed permutation tests on the product of the mean gene expression of ligand from one cell type and receptor from the other cell type. This procedure was performed between all pairs of cell types. We obtained the adjusted p-value from the permutation tests to identify the significant ligand-receptor pairs (with adjusted p-value less than 0.05, with cell type expressed proportion greater than 10%). Then the weights of the edges between each pair of cell types is determined by the number of the significant ligand-receptor pairs.

### Integration with TCGA data and survival analysis

We used the single-cell RNA-seq data to decompose the cell types of the TCGA LIHC bulk RNA-seq data using dtangle. We compared the cell type proportion between the primary solid tumor samples and normal solid tissue samples by performing Wilcoxon Rank Sum analysis. To access the relation between the proportion of cell types with survival, we applied a Cox proportional hazards model (implemented in the R package survival) that included age, gender, tumor stage in addition with the proportion of certain cell type in the sample.

## Author Contributions

JH and ZGH designed the project. JH performed all experiments and wrote the manuscript. JH, YL, QL, XS and SG analyzed the data. JY participated in discussion, language editing and reviewed the manuscript.

## Acknowledgements

This work is supported by the Project of National Science Foundation of China (NSFC 81702730) for JH.

SI-Fig 1. HE staining of the DEN-induced mice.

SI-Fig2. Gene markers of each cell types.

SI-Fig 3. Cell communication predicted by CellPhoneDB (a). The interaction between each cluster (b). Overview of selected ligand–receptor interactions; *P* values indicated by color (permutation test, see Methods).

Supplementary Information-Table 1 Parameters of each sample.

Supplementary Information-Table 2 Marker genes of the T cell subsets.

Supplementary Information-Table 3 Proportion changing Early to Late stage of subsets.

Supplementary Information-Table 4 Patterns of cell-cell communication.

