## Supplementary figures and images for "Single-Cell RNA-Seq Reveals Naïve B cells Associated with Better Prognosis of HCC"

### Supplementary Information-Fig 1

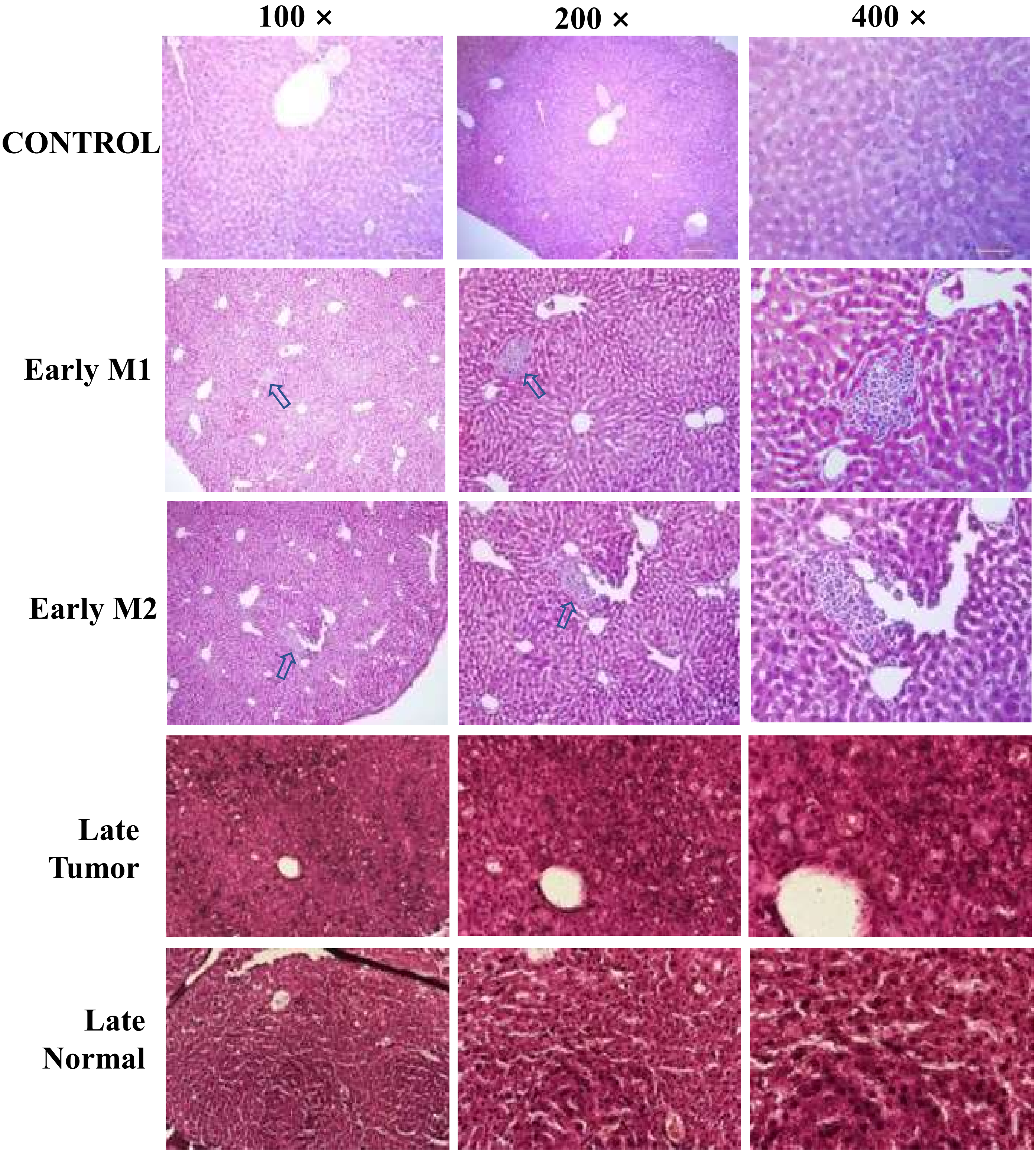


SI Fig 1. HE staining of the DEN-induced mice.

### Supplementary Information-Fig 2

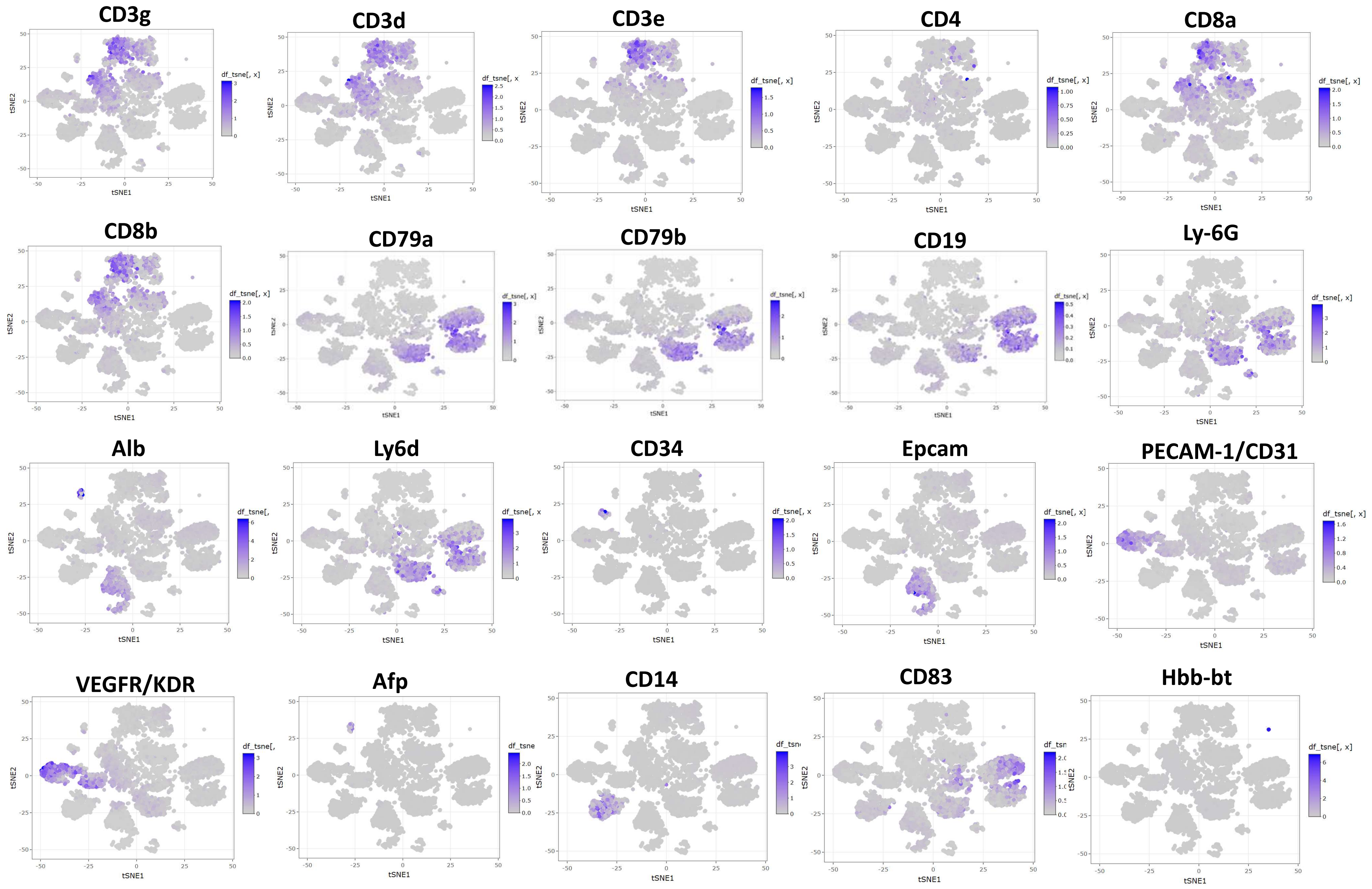


SI-Fig2. Gene markers of each cell types.
