## Supplementary Information-Fig 3 for "Single-Cell RNA-Seq Reveals Naïve B cells Associated with Better Prognosis of HCC"

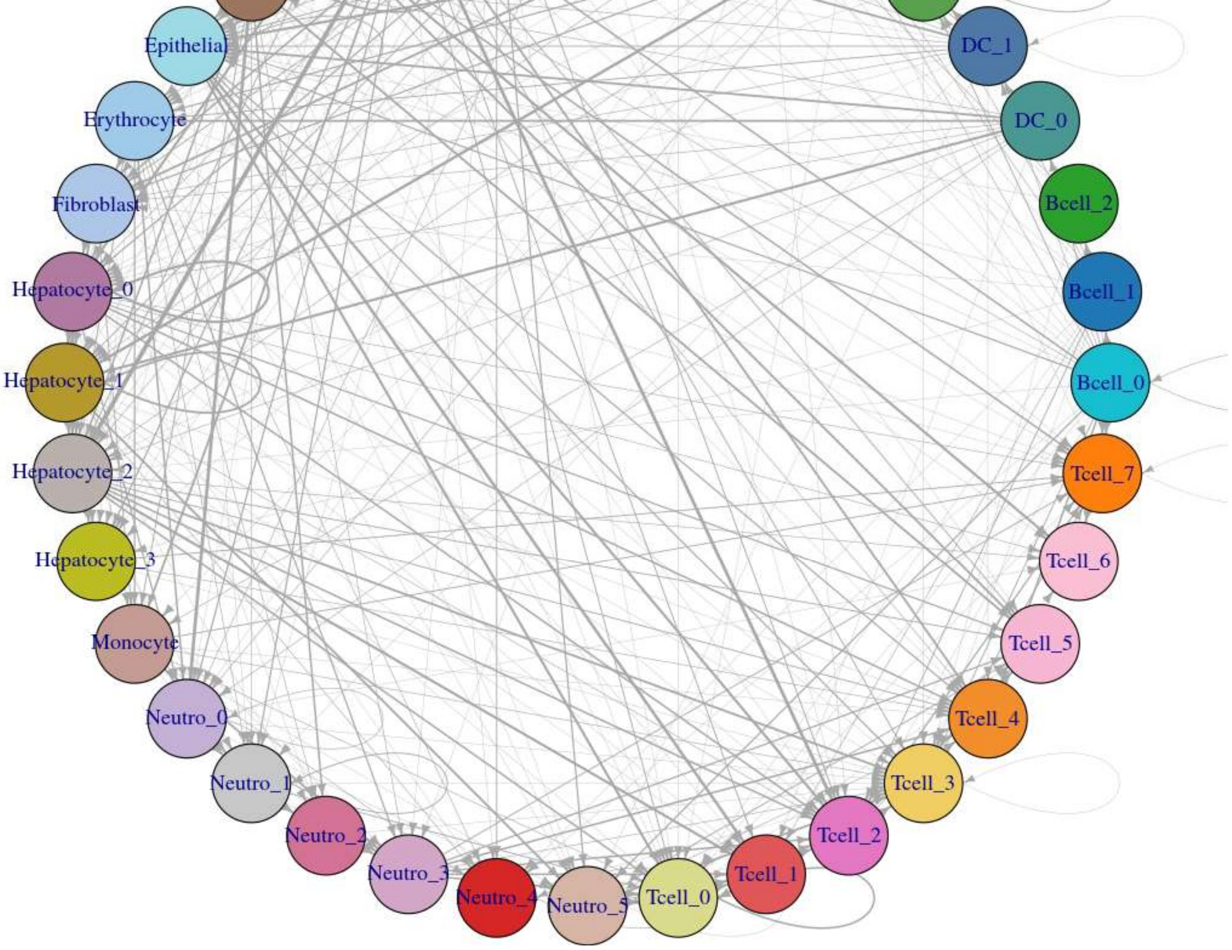
Epithelial
Erythrocyte
Fibroblast
Hepatocyte_0
Hepatocyte_1
Hepatocyte_2
Hepatocyte_3
Monocyte
Neutro_0
Neutro_1
Neutro_2
Neutro_3
Neutro_4
Neutro_5
Tcell_0
Tcell_1
Tcell_2
Tcell_3
Tcell_4
Tcell_5
Tcell_6
Tcell_7
Bcell_0
Bcell_1
Bcell_2
DC_0
DC_1

b

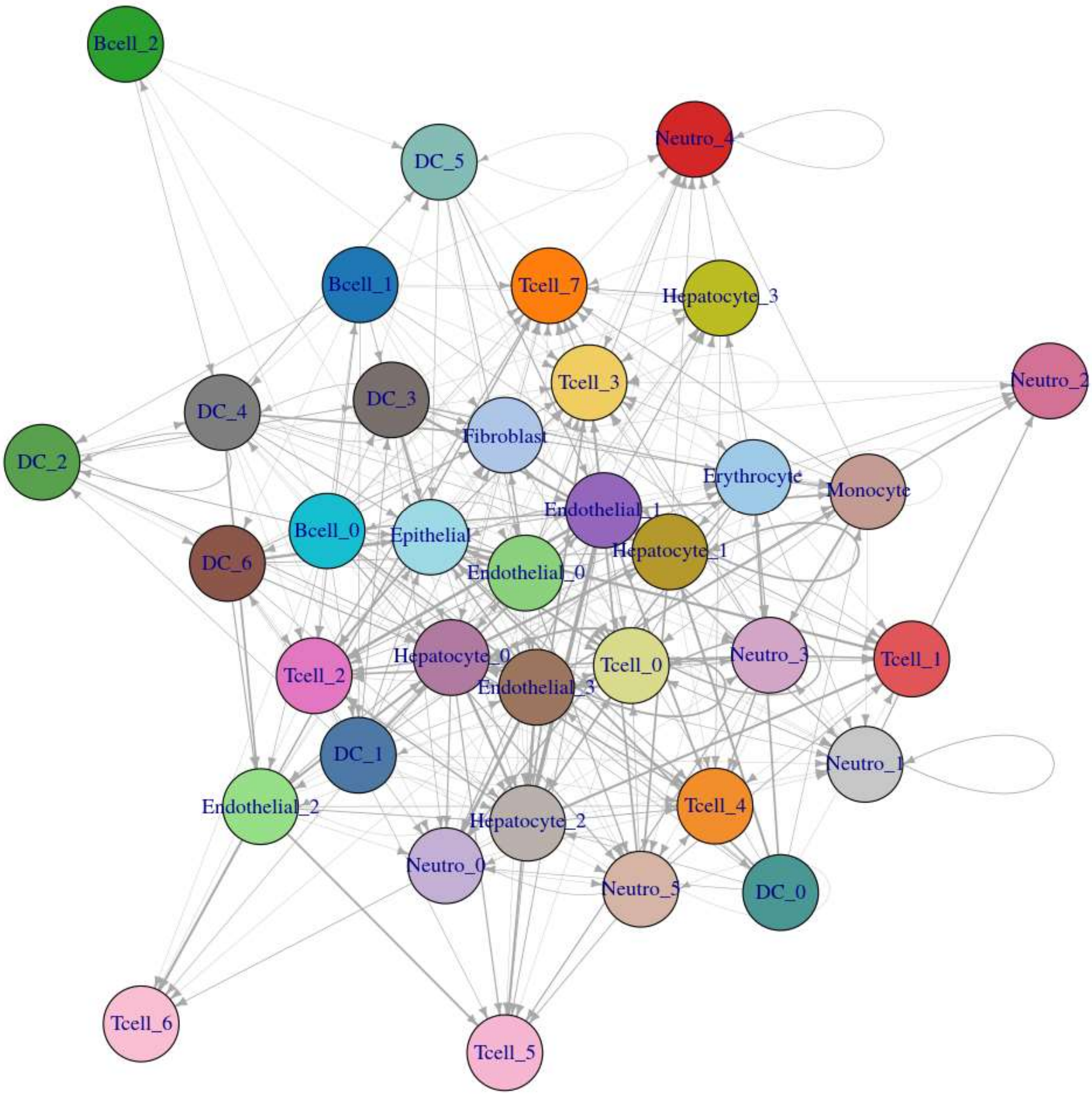
Bcell_2
DC_5
Neutro_4
Bcell_1
Tcell_7
Hepatocyte_3
Neutro_2
DC_4
DC_3
Tcell_3
DC_2
Fibroblast
Erythrocyte
Monocyte
Endothelial_1
Bcell_0
Epithelial
Hepatocyte_1
DC_6
Endothelial_0
Tcell_2
Hepatocyte_0
Endothelial_3
Tcell_0
Neutro_3
Tcell_1
DC_1
Neutro_1
Endothelial_2
Hepatocyte_2
Tcell_4
Neutro_0
Neutro_5
DC_0
Tcell_6
Tcell_5

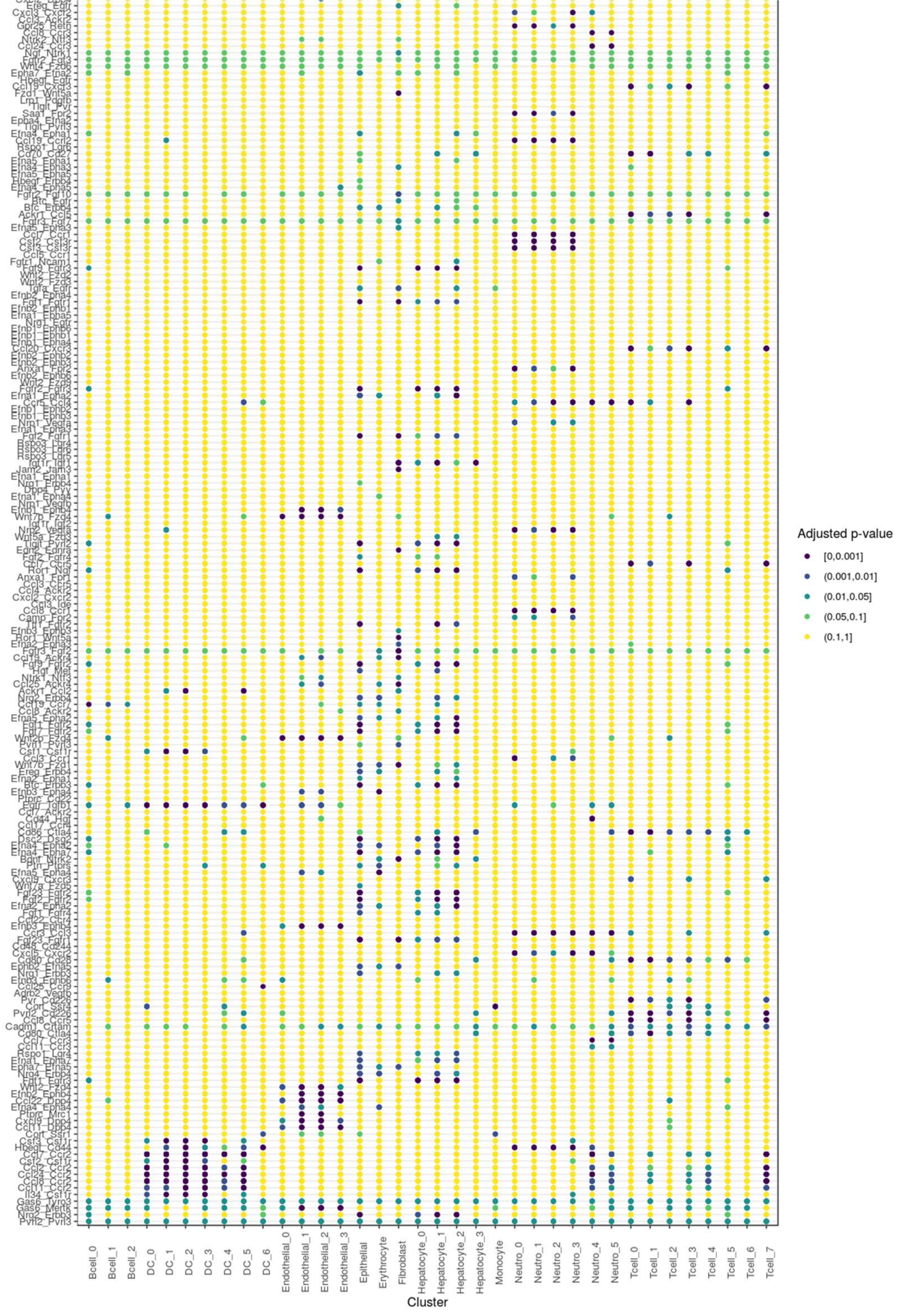


SI-Fig 3. Cell communication predicted by CellPhoneDB (a). The interaction between each cluster (b). Overview of selected ligand–
