## Supplementary Information-Table for "Single-Cell RNA-Seq Reveals Naïve B cells Associated with Better Prognosis of HCC"

Supplementary Information -Table 1 Parameters of each sample

|  | **Pre-tumor** | | **Cancer** | |
| --- | --- | --- | --- | --- |
|  | Mouse 1 | Mouse 2 | Tumor | Adjacent |
| Number of cells | 2124 | 2290 | 1572 | 1930 |
| Mean reads per cell | 102713 | 90609 | 124021 | 165148 |
| Median Genes per cell | 305 | 674 | 623 | 930 |

Supplementary Information -Table 2 Marker genes of the T cell subsets

|  | **Name** | **Description** | **Key markers** | **Notes** | **References** |
| --- | --- | --- | --- | --- | --- |
| **1** | **Tcell_0** | T0-Exhausted CD8+T cell | Lag3, CTLA3, PDCD1, HAVCR2 | Dominant in both adjacent and tumor tissue of the HCC late stage | Chen and Flies, 2013. |
| **2** | **Tcell_1** | T1_ MAIT | SLC4A10, ZBTB16, and RORC | Composed of cells from adjacent and tumor tissue of the HCC late stage/ mucosal-associated invariant T cells | Kurioka et al., 2016 |
| **3** | **Tcell_2** | T2-Cd38+T cell | Cd55, Cd38, Cd79, Cd74 | Did not show significant different between each stage |  |
| **4** | **Tcell_3** | T3-naïve T cell | LEF1, FCGR3, TCF7 | Early stage, did not show significant different between tumor and adjacent | Fo¨ rster et al., 2008 |
| **5** | **Tcell_4** | T4-NKT | GZMA, GZMK, GZMB, CXCR6 | predominantly composed of cells from early stage |  |
| **6** | **Tcell_5** | T5-Irf8+T | Irf8, Cd74 | predominantly composed of cells from adjacent and tumor tissue of the HCC late stage |  |
| **7** | **Tcell_6** | T6_Cd4+Treg cell | Cd4+, CD25 and Foxp3 | predominantly composed of cells from early stage |  |
| **8** | **Tcell_7** | T7_Effctor T | CXCCR1  FCGR3A | T cells with effector functions | Bo¨ ttcher et al., 2015 |
|  | **B cell_pan marker** | B cell | Cd79a,  Cd79b,  Cd19 |  |  |
| **1** | **B cell_0** | B0_Naïve B cell | Cd83，  Irs2 | increase in the proportion from the early to late stage tumorigenesis |  |
| **2** | **B cell_1** | B1_Plasmablast | CD27,  CD38,  SLAMF7 | dramatically decrease in the late stage |  |
| **3** | **B cell_2** | B2_Memory B | CD27,  CD38 , CD40 | dramatically decrease in the late stage |  |

Supplementary Information -Table 3 Proportion changing Early to Late stage of subsets

| Subsets | Proportion changing Early to Late stage |
| --- | --- |
| T0- Exhausted CD8+T cell | ↑ |
| T1_ MAIT | ↑ |
| T2-Cd38+T cell | -- |
| T3-naïve T cell | ↓ |
| T4-NKT | ↓ |
| T5-Irf8+T | ↑ |
| T6_Cd4+T cell | ↓ |
| T7_Effector T | ↓ |
| B0_ Naïve B cell | ↑ |
| B1_ Plasmablast | ↓ |
| B2_ Memory B | ↓ |
| DC0 | ↓ |
| DC1 | -- |
| DC2 | ↑ |
| DC3 | -- |
| DC4 | -- |
| DC5 | ↓ |
| DC6 | ↑ |
| Endothelial0 | -- |
| Endothelial1 | ↑ |
| Endothelial2 | -- |
| Endothelial3 | ↓ |
| Neutrophil0 | ↓ |
| Neutrophil1 | -- |
| Neutrophil2 | -- |
| Neutrophil3 | -- |
| Neutrophil4 | ↑ |
| Neutrophil5 | ↑ |
| Neutrophil6 | ↓ |
| Hepatocyte0 | ↓ |
| Hepatocyte1 | ↓ |
| Hepatocyte2 | ↓ |
| Hepatocyte3 | ↑ |

Supplementary Information -Table 4 Patterns of cell-cell communication

| Subsets | originating | terminal | self-regulation |
| --- | --- | --- | --- |
| T0 |  | √ |  |
| T1 |  | √ |  |
| T2 |  | √ |  |
| T3 |  | √ |  |
| T4 |  | √ |  |
| T5 |  | √ |  |
| T6 |  | √ |  |
| T7 |  | √ |  |
| B0 | √ |  | √ |
| B1 | √ |  |  |
| B2 | √ |  |  |
| Hepa0 |  |  | √ |
| Hepa1 |  |  | √ |
| Hepa2 |  | √ |  |
| Hepa3 |  | √ |  |
